# An integrated one-step assay combining thermal lysis and loop-mediated isothermal DNA amplification (LAMP) in 30 min from *E. coli* and *M. smegmatis* cells on a paper substrate

**DOI:** 10.1101/594374

**Authors:** Priyanka Naik, Siddhant Jaitpal, Prasad Shetty, Debjani Paul

## Abstract

Developing sensors in the domains of food safety, soil analysis, water quality monitoring and healthcare often requires distinguishing between different species of bacteria. The most rapid, sensitive and specific method to identify bacteria is by analysing their DNA sequence, which comprises of disinfection and lysis of bacterial cells, amplification of the isolated DNA and detection of the amplified sequence. Seamless integration of these assays on a paper substrate remains a big challenge in paperfluidic nucleic acid analyis. Combining lysis and isothermal amplification in a single reaction step is difficult because the porosity of paper and the presence of cell debris following lysis reduces the efficiency of DNA amplification. On the other hand, extracting and purifying the DNA after lysis to improve the amplification efficiency involves addition of chemical reagents, one or more wash steps and manual intervention. This problem is even more complex for mycobacteria as its thick cell wall structure impedes lysis and the high GC-content of the genome requires careful optimization of enzymatic denaturation during isothermal amplification. Here we successfully combine thermal lysis and loop-mediated isothermal amplification (LAMP) into a single reaction step on paper without the need for any intermediate intervention. We demonstrate our integrated assay by amplifying DNA from 100 CFU/mL of *Escherichia coli* (MG1655) and *Mycobacterium smegmatis* (mc^2^155) cells in 30 min on a paper substrate. We also confirm that *E. coli* and *M. smegmatis* can be completely disinfected on paper by heating at 60 °C for 5 min and 15 min respectively, making this assay safe and suitable for incorporation into diverse paperfluidic sensors for field use.

Electronic Supplementary Information (ESI) is available.

## 1. Introduction

Bacterial sensors have applications in diverse fields ranging from clinical diagnostics to food safety, water analysis and environmental monitoring ^1^. Conventional assays to distinguish different species of bacteria are time-consuming (e.g. cultures) or prone to false positives (e.g. antigen-based tests) to be effectively incorporated into sensors ^2,3^. DNA analysis, on the other hand, is a rapid, sensitive and specific technique for identifying different bacteria. Varadi *et al.* reviewed the various methods for detection and identification of bacteria, one of which was DNA analysis. Their report highlights the strength of nucleic acid amplification to generate enhanced signals from complex samples containing low numbers of bacteria of interest amidst billions of other cells. This facilitates the identification of specific virulence or resistance mechanisms which could be useful in administering pathogen-directed clinical treatment and address the growing problem of antimicrobial resistance. The authors also point out that DNA analysis allows rapid detection of bacteria which are slow growing or difficult to culture e.g. *Mycobacterium* spp ^4^.

Paper has emerged as a popular substrate for developing affordable sensors for nucleic acid analysis, which comprises of three main steps: (a) cell lysis, (b) amplification of one or more specific sequences in the extracted DNA, and (c) detection of the amplified sequences. While each of the three steps of DNA analysis have been individually demonstrated for detecting bacteria on paper substrates, integrating them onto a single paperfluidic device has been a major challenge. There are several examples of integrated paperfluidic devices for DNA amplification and detection, the last two steps in nucleic acid analysis ^5–8^. For example, Linnes *et al.* amplified *Chlamydia trachomatis* on chromatography paper followed by lateral-flow assay-based detection on nitrocellulose membrane ^5^. Tsaloglou *et al*. performed DNA amplification and electrochemical detection on a paper substrate with purified genomic DNA of *M. smegmatis* and *M. tuberculosis* ^8^. Fu *et al.* developed an integrated device for PCR of *L. monocytogenes* DNA, followed by lateral flow detection of the amplified DNA on paper ^9^. It should be noted that DNA amplification in this case was not performed on a paper substrate. Combining sample preparation (cell lysis) and DNA amplification has proven to be much more difficult. This is because the porous network of paper and the cell debris left behind after lysis reduces the efficiency of DNA amplification. In their current formats, nucleic acid extraction techniques are not suitable for integration into a POC device. This is because these are laborious, time-consuming, and often require specialized equipment ^10^. Another caveat is the lack of efficiency and reliability since the entire process might end up as “garbage-in, garbage out” if the sample obtained is impure ^11^. Most of the studies reported in literature that have managed to perform cell lysis on paper substrates use chemical lysis reagents such as, surfactants or guanidinium-based chemicals, which are known to interfere with subsequent DNA amplification ^12^.

Isothermal amplification techniques such as helicase dependent amplification (HDA), loop-mediated DNA amplification (LAMP), recombinase polymerase amplification (RPA), nucleic acid sequence-based amplification (NASBA) have emerged as viable alternatives to PCR as the technique to amplify DNA in paperfluidic devices ^13^. These techniques avoid thermal cycling and the use of expensive equipment ($2000 - $4000). Since isothermal techniques function at a single temperature, they can be performed with inexpensive heat sources ($40 - $100) such as, hot plate, heat blocks and hand/toe warmers ^14–16^. LAMP is one of the most popular isothermal amplification methods for the development of integrated paperfluidic devices because it is extremely sensitive, specific and tolerant to several amplification inhibitors ^17^. It takes place at a single temperature between 60 °C and 65 °C and generates target amplicons of different sizes in a single reaction step ^18^. LAMP has been used to detect *S. aureus, N meningitides, S. pneumoniae and H. influenzae* in a PDMS/paper hybrid device, and for multiplexed detection of *Staphylococcus* and *Streptococcus* species on a paperfluidic device ^19–21^.

In all of these reports ^5,8,9,19–21^, purified nucleic acids were used as the starting material implying off-chip sample preparation. Sample preparation requires 1 - 2 h of work prior to the actual amplification with the additional requirement of trained personnel for sample handling. Sample preparation on paper is the most cumbersome module in the amplification of nucleic acids when the starting material is unprocessed cells. Firstly, the lysates generated after cell lysis might interfere with nucleic acid amplification. This can be avoided by either purifying the DNA before amplification or using an amplification technique that is insensitive to the presence of cellular debris. Secondly, the background signal from lysates should be much lower compared to the signal obtained from the amplified DNA. Recently several reports of complete DNA analysis on paper have emerged for detection of gram negative and gram positive bacteria. Lafleur *et al.* combined cellulose sheets and glass fibre substrates to develop a prototype for combined lysis, amplification and detection using strand displacement amplification (SDA) of methicillin-resistant *Staphylococcus aureus* ^22^. Horst *et al.* developed a prototype which combined lysis, HDA and detection on polyether sulphone (PES) membrane for diagnosis of gonorrhoea ^16^. Another study by Tang *et al*. reports an integrated device employing HDA for detection of *S. typhimurium* in various spiked samples ^23^. The residual lysis and the precipitation buffer components used in these studies are likely to have reduced the sensitivity of nucleic acid amplification ^24,25^. While there are various reports of integrated assays on paper (Table 1), due to extensive washing steps to filter amplification inhibitors there is an increase in the total assay time and complexity, which limits the translation of these assays. These studies suggest that the development of integrated devices is impeded by the tedious unification of all the modules associated with DNA analysis.

**Table 1:**
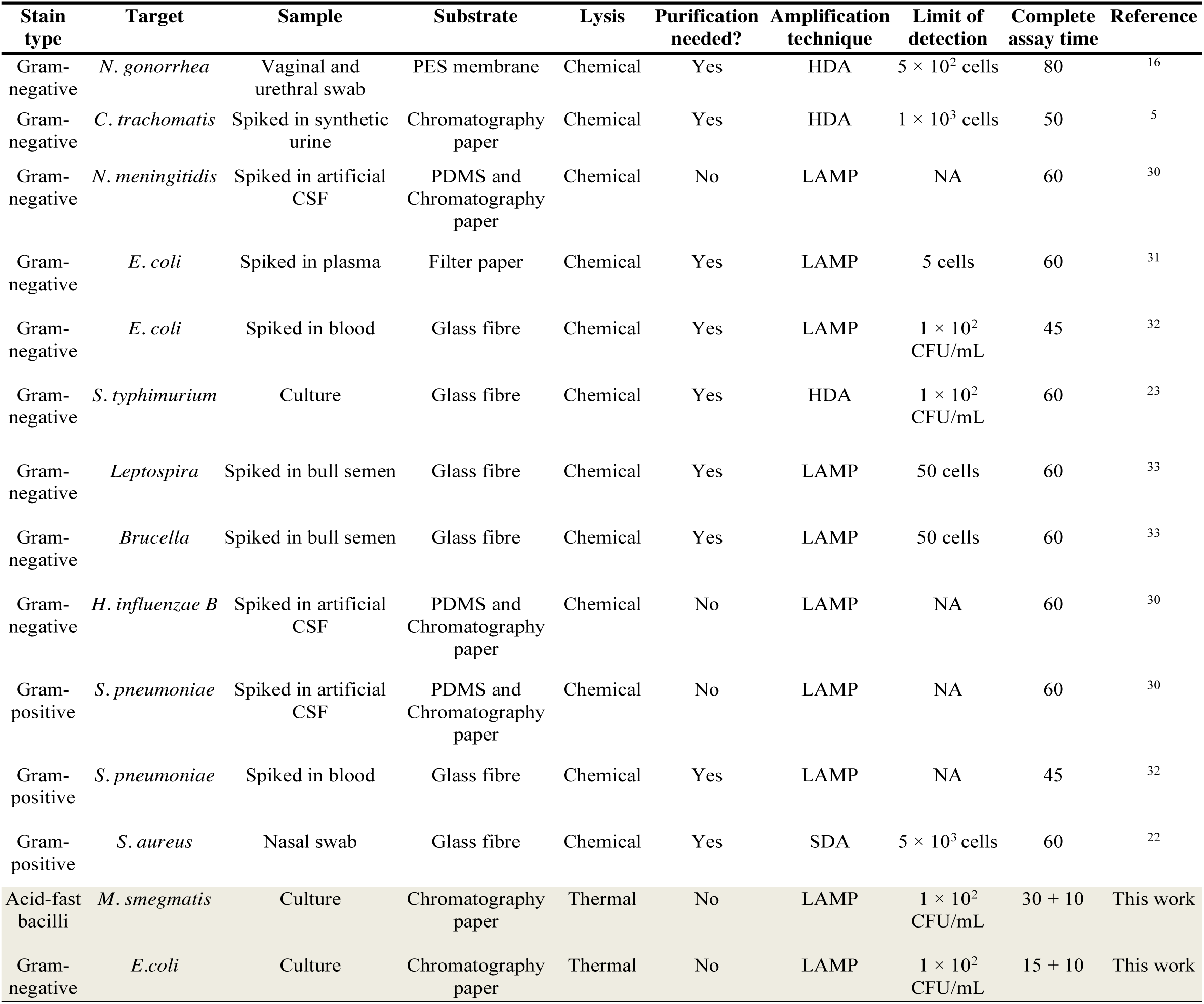
Comparison of paper-based integrated assays for detection of bacteria

| Stain type | Target | Sample | Substrate | Lysis | Purification needed? | Amplification technique | Limit of detection | Complete assay time | Reference |
| --- | --- | --- | --- | --- | --- | --- | --- | --- | --- |
| Gram-negative | *N. gonorrhea* | Vaginal and urethral swab | PES membrane | Chemical | Yes | HDA | $5 \times 10^2$ cells | 80 | 16 |
| Gram-negative | *C. trachomatis* | Spiked in synthetic urine | Chromatography paper | Chemical | Yes | HDA | $1 \times 10^3$ cells | 50 | 5 |
| Gram-negative | *N. meningitidis* | Spiked in artificial CSF | PDMS and Chromatography paper | Chemical | No | LAMP | NA | 60 | 30 |
| Gram-negative | *E. coli* | Spiked in plasma | Filter paper | Chemical | Yes | LAMP | 5 cells | 60 | 31 |
| Gram-negative | *E. coli* | Spiked in blood | Glass fibre | Chemical | Yes | LAMP | $1 \times 10^2$ CFU/mL | 45 | 32 |
| Gram-negative | *S. typhimurium* | Culture | Glass fibre | Chemical | Yes | HDA | $1 \times 10^2$ CFU/mL | 60 | 23 |
| Gram-negative | *Leptospira* | Spiked in bull semen | Glass fibre | Chemical | Yes | LAMP | 50 cells | 60 | 33 |
| Gram-negative | *Brucella* | Spiked in bull semen | Glass fibre | Chemical | Yes | LAMP | 50 cells | 60 | 33 |
| Gram-negative | *H. influenzae B* | Spiked in artificial CSF | PDMS and Chromatography paper | Chemical | No | LAMP | NA | 60 | 30 |
| Gram-positive | *S. pneumoniae* | Spiked in artificial CSF | PDMS and Chromatography paper | Chemical | No | LAMP | NA | 60 | 30 |
| Gram-positive | *S. pneumoniae* | Spiked in blood | Glass fibre | Chemical | Yes | LAMP | NA | 45 | 32 |
| Gram-positive | *S. aureus* | Nasal swab | Glass fibre | Chemical | Yes | SDA | $5 \times 10^3$ cells | 60 | 22 |
| Acid-fast bacilli | *M. smegmatis* | Culture | Chromatography paper | Thermal | No | LAMP | $1 \times 10^2$ CFU/mL | 30 + 10 | This work |
| Gram-negative | *E. coli* | Culture | Chromatography paper | Thermal | No | LAMP | $1 \times 10^2$ CFU/mL | 15 + 10 | This work |

Combining lysis and amplification into a single reaction on paper is especially challenging for mycobacteria because they have a thick and waxy cell wall structure due to the presence mycolic acid ^26^. Also, the mycobacteria genome has a high GC content which impedes the enzymatic denaturation process employed in isothermal amplification techniques. We have earlier demonstrated one-step lysis and amplification using HDA at 65 °C of *Mycobacterium tuberculosis* cells in a solution-based reaction ^14^. We have also demonstrated amplification of an 84 bp purified DNA fragment of *M. tuberculosis* on a paper substrate ^15^. There are no reports of successful amplification of DNA from unprocessed mycobacteria cells on a paper substrate. In this study, we have combined disinfection, thermal lysis and LAMP directly into a single reaction step on paper without any wash, purification or manual intervention. We demonstrated this assay on 100 CFU/mL of unprocessed *Mycobacterium smegmatis* (mc^2^155) (a surrogate for *M. tuberculosis*) and *E. coli* (MG1655) cells in 30 min. In our protocol, *E. coli* and *M. smegmatis* are disinfected on paper by heating at 60 °C for 5 min and 15 min respectively. We believe the integration of sample preparation with DNA amplification into a single step makes our assay ideal for incorporation into different kinds of paperfluidic bacterial sensors.

## 2. Materials and methods

### 2.1 Equipment and chemicals

The bacterial cultures were incubated in incubator purchased from Pooja Lab Equipment (Mumbai, India). The optical density of the bacterial cultures was measured using a spectrophotometer (Spectroquant Pharo 100, Merck Millipore, India). A 450 W heat sealer (PFS-300P, Impulse, India) was used to heat seal the paper substrate in a plastic pouch. A thermal cycler (MJ mini, Bio-Rad, India) and a hot plate (Spin Not Digital MC02, Tarsons, India) were used as heating sources in the one-step reaction. A gel electrophoresis unit was bought from Genetix (Mumbai, India). Agarose gels were imaged by a gel-doc imaging system (Geliance 1000, PerkinElmer, USA) imaging system. Fluorescence intensities of the DNA amplified on paper were measured using a microplate reader (SpectraMax M2E, Molecular devices, USA).

Bovine serum albumin, Middlebrook 7H9 and albumin dextrose catalase (ADC) for growing mycobacteria were purchased from HiMedia (Mumbai, India). Middlebrook 7H11 agar base, Middlebrook oleic albumin dextrose catalase (OADC) enrichment medium, Luria-Bertani agar base, Luria-Bertani broth base, and glycerol were purchased from Sigma Aldrich (Mumbai, India). *Escherichia coli* strain MG1655 (CGSC 6300) was procured from *E. coli* Genetic Stock Center (Connecticut, USA). DNA-Exitus plus was bought from PlanetScience (Mumbai, India). Filter pipette tips were purchased from Biotix Inc (San Diego, USA). WarmStart LAMP Kit (DNA and RNA) enzyme mix for the LAMP reactions was bought from New England Biolabs (Beverly, USA). LAMP primers were synthesized by Integrated DNA Technology (Iowa, USA) and Eurofins (Luxembourg). DNAse-free water was purchased from GeNei (Bangalore, India). Whatman Grade 1 chromatography paper for the paper-based reactions was obtained from GE Healthcare (Mumbai, India). DNA ladder (O’GeneRuler ultra low range 10–300 bp ladder), Quant-iT PicoGreen dsDNA reagent, and ethidium bromide were procured from Thermo Scientific (Mumbai, India). Agarose was purchased from Merck (Mumbai, India). Fluorescence imaging was performed in 96-well black clear bottom plates from Corning Costar (Mumbai, India).

### 2.2 Culturing of bacteria

*Escherichia coli* (MG1655) was plated on LB agar plates. Single colonies from the culture plate were transferred in LB broth. The broth was prepared by mixing 2 g of LB broth base with 100 mL of water. It was autoclaved at 121 °C for 20 min, cooled down to 40 °C followed by addition of 1% inoculum to the broth. The inoculated broth was incubated overnight at 37 °C and 240 rpm. The bacteria were diluted using LB broth to concentrations ranging from 1–10^7^ CFU/mL for subsequent experiments.

*Mycobacterium smegmatis* (mc^2^155) was plated on M7H11 agar plates supplemented with Middlebrook ADC growth supplement. Single colonies selected from the plates were grown in M7H9 broth. The broth was prepared by mixing 2.35 g M7H9 and 2 mL of glycerol in 450 mL of water. It was autoclaved at 121 °C for 20 min. Middlebrook OADC growth supplement was added aseptically to the broth after it cooled down to 40 °C. Inoculum (1%) was added to the broth followed by incubation at 37 °C and 240 rpm for 15 h. The bacteria were diluted using M7H9 broth to concentrations in the range of 1 – 10^8^ CFU/mL for subsequent experiments.

### 2.3 Thermal lysis of bacteria

Cell viability after thermal lysis at 60 °C was assessed by spotting 2 μL of *E. coli* on 5 mm diameter paper discs. Each paper disc was then heat-sealed in a polythene pouch which was subjected to 60 °C for 5 min, 10 min, 15 min, 30 min, 45 min, and 60 min respectively. Each paper disc was held using forceps and streaked on LB agar plate to distribute the bacteria across the agar plate. The lysis experiments were performed in triplicates. The plates were then incubated at 37 °C overnight and imaged to see any visible bacterial colonies. Cell viability of *M. smegmatis* was also assessed in a similar manner. M7H11 agar plates were used with incubation at 37 °C for 72 h.

### 2.4 DNA amplification from unprocessed cells on paper

A 10 µL LAMP reaction mixture consisted of 5 µL of the WarmStart LAMP 2X master-mix, and 2 µL of the template (50 ng DNA or cells). LAMP primers for *M. smegmatis* and *E. coli* were the same as described by Iwamoto *et al.* and Stratakos *et al.* ^27,28^. The paper was pre-treated as described in our earlier work ^15^. Appropriate negative and positive controls were set up during each experiment. The negative controls consisted of the no master-mix control (NMC) and no template control (NTC). NMC consisted of DNase free water, primers and the cells. NTC contained everything else except the cells. As shown in figure 1, the reaction mixture containing bacterial culture was added to a 5 mm diameter paper disc, heat-sealed into a plastic pouch and incubated at 60 °C for varying durations.

**Figure 1.**
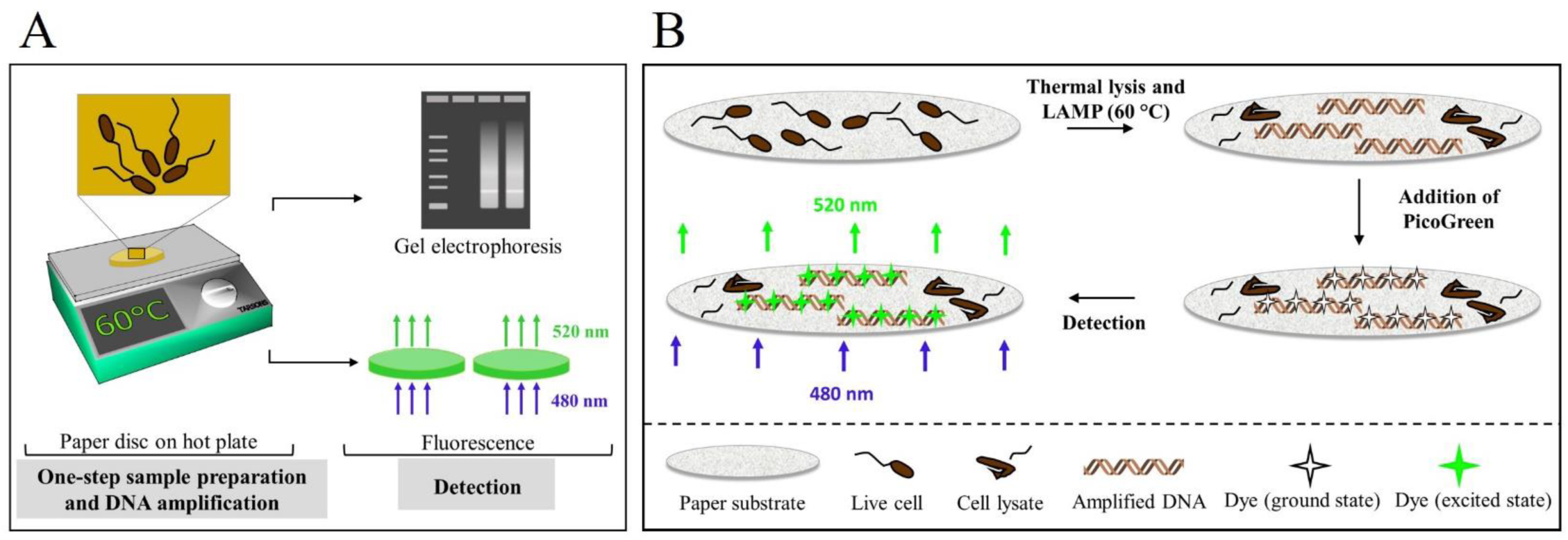
Panel A shows the schematic of the one-step lysis and LAMP protocol. The LAMP reaction mix is spotted on a pre-treated 5 mm paper disc. The paper disc is put in a polyethylene pouch, which is then heat sealed. The pouch is then subjected to a temperature of 60 °C for LAMP. PicoGreen is spotted on the paper disc after amplification and allowed to bind to the dsDNA for 5 min. Fluorescence intensity (Ex./Em.: 480 nm/ 520 nm) is measured using the microplate reader. Panel B shows the schematic diagram of the biochemistry of the reaction. The thermal lysis causes release of the genomic DNA which serves as the template for subsequent DNA amplification. PicoGreen is added to the mixture which binds to the dsDNA molecules. The recorded fluorescence intensities were then analysed to confirm the efficacy of the one-step protocol.

### 2.5 Detection of the amplified DNA

Amplified DNA was detected by electrophoresis using a 2% agarose gel containing 0.5 µg/mL ethidium bromide. Paper discs containing the amplified product were directly loaded in the wells of the gel using forceps. The DNA bands were observed under the UV gel doc system.

We also detected the amplified DNA using fluorescence. A 96-well black clear-bottom plate was modified using PDMS before detecting the fluorescence signal from the paper substrate using a plate reader. PDMS base and curing agent were mixed in the ratio of 10:1, degassed, poured in the wells up to 5 mm height, and cured at 65 °C for 1 h. This strategy was adopted as the microplate reader could not reliably track fluorescence emitted by the PicoGreen-soaked paper discs when placed in the bottom of the well (i.e. in the absence of PDMS). PicoGreen (1:10 final dilution) was spotted on the paper disc after the amplification and was incubated at room temperature for 5 min. The paper discs were subjected to a top read using excitation/emission wavelengths of 480 nm/520 nm in the microplate reader.

### 2.6 Statistical analysis

One-way ANOVA with Tukey’s post-hoc test was performed to compare the data amongst the control and the test groups. Data were expressed as mean ± standard deviation of replicates of independent experiments. A p-value < 0.05 was reported as statistically significant.

## 3. Results and discussion

### 3.1 We can thermally lyse bacteria on a paper substrate at 60°C in 5 min

We developed a protocol that completely kills bacteria on paper and amplifies the released DNA at 60 °C. Our initial studies were done using *E. coli* MG1655. *E. coli* is a common inhabitant in a variety of specimens from soil and vegetables, raw meat, water to mammalian gastro-intestinal tract. Our protocol efficiently lyses *E. coli* in 5 min (figure 2, top panel). We did not explore the efficacy of lysis at lower temperatures because the manufacturer of the LAMP kit recommends an operating temperature range of 60 °C – 65 °C. We further tested the disinfection efficiency of *M. smegmatis*. As shown in figure 2, the unlysed plate (negative control) shows abundant bacterial growth. As expected, lysis at the recommended temperature of 95 °C for 30 min kills the bacteria completely and hence serves as a positive control. We varied lysis times from 60 min to 5 min. We find that thermal lysis for 15 min is sufficient to kill all mycobacteria as the plates did not show any visible growth even after 7 days of incubation. However, as seen in figure S1, lysis times less than 15 min led to growth of bacteria.

**Figure 2.**
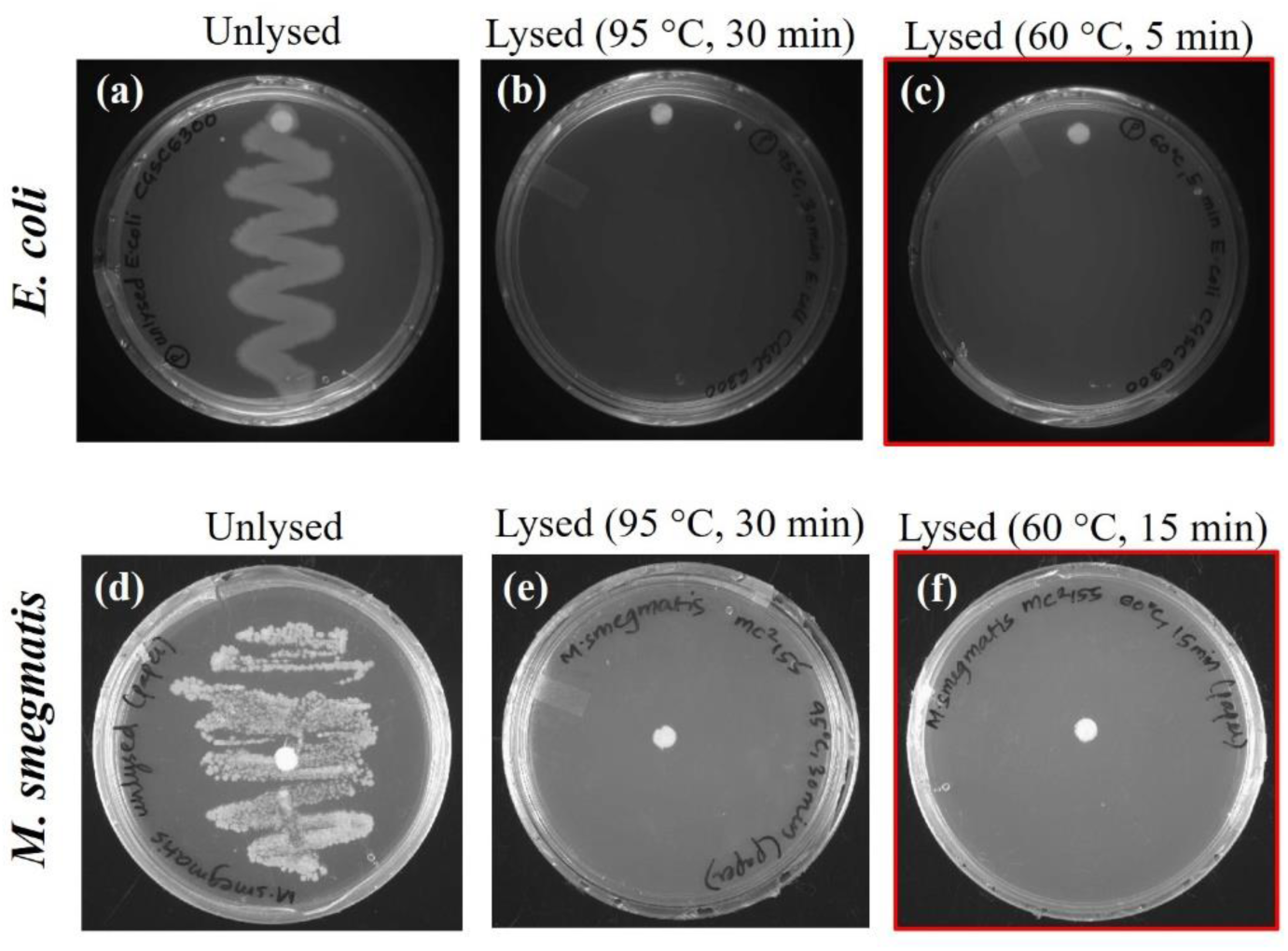
Disinfection following thermal lysis of *E*. coli MG1655 (top panel) and *M. smegmatis* mc^2^155 (bottom panel). Panels (a, d) show the plate with unlysed cells (negative control). Panels (b, e) indicates the plate with lysis on paper at the recommended protocol of 95 °C for 30 min (positive control). Panels (c, f) show the result of thermal lysis on paper at a lower temperature (60 °C) and a reduced duration (5 min and 15 min). The images were taken after incubation at 37 °C for 24 h for *E. coli* and 72 h for *M. smegmatis.* The white spots are the paper discs. All experiments were performed in triplicates.

### 3.2 How to prevent carryover contamination when performing LAMP on paper substrates?

LAMP is an extremely sensitive technique and it is reported to be highly prone to cross-contamination ^29^, as also witnessed by us during the course of our study. All of the reported measures to prevent amplification in the negative controls focus on solution-based reactions carried out in PCR tubes. The most critical piece of advice in this case is not to open the tubes after the reaction. In our case, the paper substrate often needed to be taken out for loading in the gel or to measure fluorescence. Therefore, we had to develop our own set of measures to manage cross-contamination. The most useful step we found was wiping the lab bench using DNA Exitus every time before setting up a reaction. The next most important step was to have four separate work areas for each step of the process: (1) preparation of the primer solutions, (2) addition of the reaction mixture to the paper discs, (3) combined thermal lysis and amplification on the hot plate, and (4) detection of the amplified DNA using gel electrophoresis or fluorescence. Additionally, we autoclaved the pipettes and forceps before each reaction, and used facemasks while pipetting the reaction mixtures on paper.

### 3.3 We can amplify and detect DNA from unprocessed *E. coli* (MG1655) and *M. smegmatis* (mc^2^155) cells in 30 min

Once we ensured that thermal lysis at 60 °C is effective, we wanted to combine it with LAMP on the same piece of paper without any intermediate purification of the extracted DNA. We varied the total reaction times (lysis + amplification) from 45 min to 5 min for both *E. coli* MG1655 (10^7^ CFU/mL) and *M. smegmatis* mc^2^155 (10^8^ CFU/mL). We used the highest concentration of the bacteria for this experiment as we wished to determine the minimum time required for us to get a significant fluorescence signal compared to the negative control.

We detected the amplified DNA by fluorescence from the DNA binding dye PicoGreen as it allowed us to easily quantify the assay efficiency. We added PicoGreen to the paper substrate after the amplification step because we observed a significant decrease in its fluorescence intensity when the dye is heated (figure S2). Shetty *et al*. explored different DNA-binding dyes and recommended PicoGreen as the most suitable dye for detecting amplified DNA on a paper substrate ^15^. They did not have cell debris in their reaction mixture as they used externally purified DNA as their template. Therefore, we first confirmed that PicoGreen shows negligible fluorescence on a paper substrate in presence of cell debris (data not shown). Amplified DNA from *E. coli* and *M. smegmatis* could be detected in 15 min and 30 min respectively with an additional 10 min to obtain the fluorescent readout (figure 3A and B). The longer reaction time for *M. smegmatis* is expected due to the differences in the cell wall structure and the GC content of the genome of the two bacteria. The difference in fluorescence between the different one-step test reactions and the negative controls (e.g. no mastermix and no template) was significant (p-value < 0.05). The test reactions showed no statistically significant difference when compared with the positive control (60 min), as shown in figure 3A and B. Gel electrophoresis was performed as an additional check to confirm amplification on the paper substrate (figure S3).

**Figure 3.**
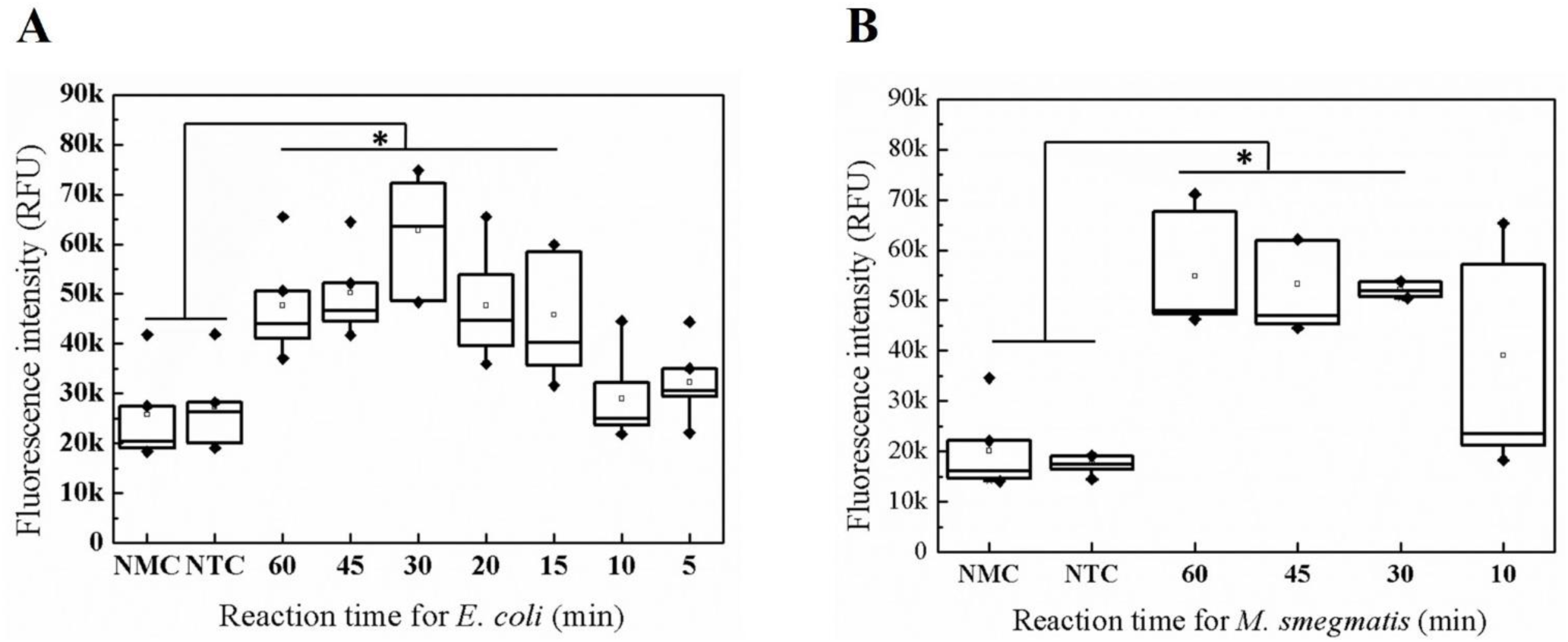
One-step lysis and direct DNA amplification from *E. coli* (Panel A) and *M. smegmatis* (Panel B) on paper. Panels (A, B) show at what minimum reaction time the amplified DNA (n = 6) can be detected over the background. The reactions for *E. coli* were performed for 5 min, 10 min, 15 min, 20 min, 30 min, 45 min and 60 min. The reactions for *M. smegmatis* were performed for 30 min, 45 min and 60 min. A p-value < 0.05 confirms statistically significant difference between the negative controls (NMC and NTC) and the experimental groups. No significant difference was seen between the experimental groups and the positive control (60 min). According to the fluorescence readouts, the minimum time taken for combined lysis and amplification on paper for *E. coli* and *M. smegmatis* is 15 min and 30 min respectively.

### 3.4 We can detect 100 CFU/mL of *E. coli* (MG1655) and *M. smegmatis* (mc^2^155) using our one-step assay

We next tested the sensitivity of our combined reaction protocol on several different concentrations of *E. coli* and *M. smegmatis*. For this assay, we used the minimum reaction times obtained from our previous study (i.e. 15 min for *E. coli* and 30 min for *M. smegmatis*). The detection sensitivity of our protocol was 100 CFU/mL for both the bacteria when detected using fluorescence (figure 4A and B). The limit of detection of the amplified products was also tested by gel electrophoresis (figure S4). As seen in table 1, sensitivity of our assay is comparable to the other integrated assays on paper that have been reported in the literature.

**Figure 4.**
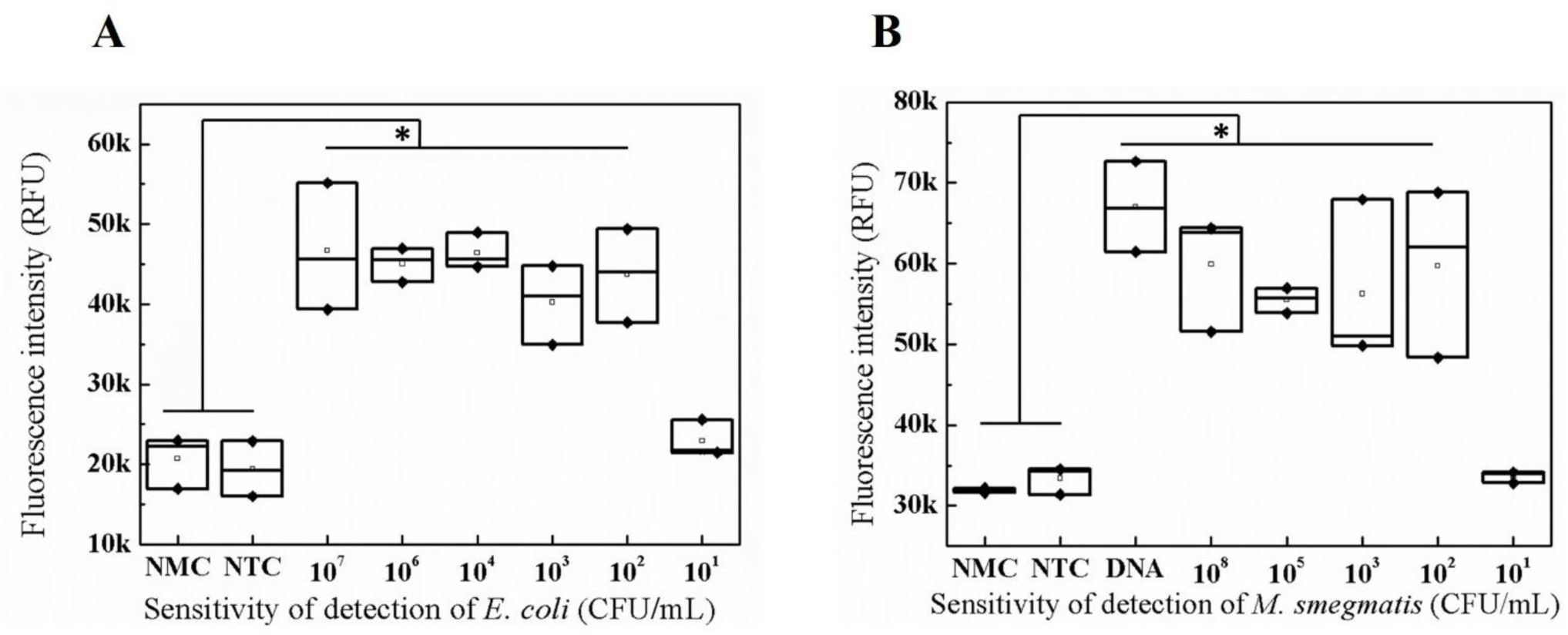
The detection sensitivity of our integrated assay (n = 3) with *E. coli* (panel A) and *M. smegmatis* (panel B). A p-value < 0.05 confirms statistically significant difference between the negative controls (NMC and NTC) and the experimental groups. No significant difference was seen between the experimental groups and the positive control (purified DNA). According to the fluorescence data, the limit of detection for *E. coli* and *M. smegmatis* is 100 CFU/mL.

### 3.5 Comparison of our assay with other integrated tests in the literature

We compared our assay with different paper-based integrated devices (i.e. combined sample preparation, DNA amplification and detection) reported in the literature. More specifically, we compared the nature of the sample (clinical or spiked), reaction substrate, lysis technique, the method used for DNA amplification, whether any purification is needed between lysis and amplification steps, the limit of detection, and the time taken to complete the entire assay from start to finish. The table shows that an overwhelming majority of the reported assays involves detection of gram-negative bacteria. This is because their thin lipopolysaccharide cell wall structure allows for facile lysis, making it easy to adapt these assays to paperfluidics. This observation matches our findings with *E. coli*.

All the reports summarized in table 1 used chemical lysis to disrupt the bacterial cell wall. Most of them also required intermediate intervention, typically in the form of a wash step, between lysis and amplification steps to get rid of the inhibitory reagents. Dou and others allowed the solvents used in lysis buffer to evaporate instead of performing a wash, and then added the amplification mixture to the paper ^30^. We employed thermal lysis which does not require any additional reagents, thereby obviating the need for any wash step. We can load bacteria directly onto the paper substrate, add the amplification mixture, and perform both lysis and amplification in the same step by heating it at 60 °C for 30 min. To the best of our knowledge, this is the first report which demonstrates combined thermal lysis and amplification on a paper substrate in a single uninterrupted step.

As expected, most of these assays have employed LAMP to amplify DNA due to its high sensitivity and specificity. Glass fibre and paper substrates (chromatography or filter) seem to be the most popular choices for nucleic acid amplification assays in general. For ease of comparison, we converted the limits of detection (LOD) to cells/mL in some of the cases by considering the reported reaction volumes. Values reported in CFU/mL are presented as such. We did not include those reports which used externally purified nucleic acids as the starting material. Our detection limit of 100 CFU/mL compares favourably with most of the integrated assays summarized in table 1.

We finally compared the total assay time of various reports by considering the time taken from adding the sample containing bacteria to the substrate until it is detected. Our total assay time of 40 min is similar to that of other assays. Overall, our assay on paper compares very favourably to the integrated assays reported on paper, with the additional advantage of it being a single step reaction with a minimum number of reagents.

## 4. Conclusions

We have successfully demonstrated one-step disinfection, thermal lysis and DNA amplification of 100 CFU/mL of *Escherichia* coli and *Mycobacterium smegmatis* on paper in 30 min. We have detected the amplicons on paper using the fluorescence from the DNA-binding dye PicoGreen. We showed that thermal lysis at 60 °C effectively kills all bacteria, making this assay safe and suitable for incorporation into sensors. Our detection sensitivity compares well to the reported integrated paper-based nucleic acid amplification techniques. Our next immediate goal is to validate our assay with complex clinical samples. We also plan to incorporate our one-step protocol into a device for biosensing at the point of care.

## 6. Acknowledgements

We acknowledge funding from the Infrastructure Facility for Advanced Research and Education in Diagnostics, Dept. of Biotechnology and Wadhwani Research Centre for Bioengineering, IIT Bombay (BT/INF/22/SP23026/2017). PN acknowledges the WHEELS Global Foundation (RD/0117-DON00G0-001 and DO/2017-SUMP001-001) for salary support and project expenses. SJ acknowledges Wadhwani Research Centre for Bioengineering, IIT Bombay (RD/0116-DONW0G0-010) for salary support. We thank Dr. Sarika Mehra (IIT Bombay) for providing the *M. smegmatis* (mc^2^155) strain. We also thank Bhavik Shah, Amrita Sharma, Rakesh Kumar and Rishu Tiwari for use of their gel doc system and microplate reader. We thank Srushti Singh, Md. Ramiz Raza, Shilpi Pandey, Riddha Mana and Santosh Jinnawar for technical assistance and feedback on the manuscript.

## Supplementary information

### 1. Thermal lysis of *M. smegmatis* for durations under 15 min is inefficient

Cell viability of *M. smegmatis* was assessed after thermal lysis at 60 °C by spotting 2 µL of bacteria on 5 mm diameter paper discs. Each paper disc was then heat-sealed inside a polythene pouch and heated at 60 °C for 5 min and 10 min respectively. The pouches were cut open and the paper discs were streaked on M7H11 agar plates. These plates were incubated at 37 °C for 72 h to monitor the bacterial growth after lysis. Unlysed bacteria were used as a negative control, while bacteria thermally lysed for 30 min at 95 °C were used as a positive control. As seen from the images of the plates in figure S1, heating *M. smegmatis* for durations under 15 min does not lead to complete disinfection.

**Figure S1.**
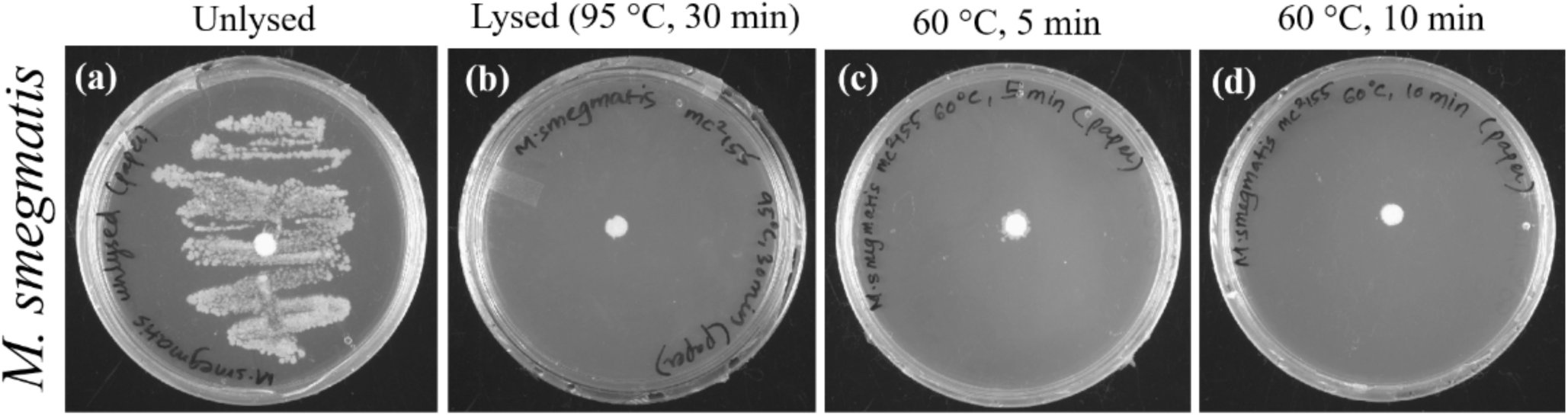
Disinfection and thermal lysis of *M smegmatis* (mc^2^155). Panel (a) shows the plate with unlysed cells (negative control). Panel (b) indicates the plate with lysis on paper at the recommended protocol of 95 °C for 30 min (positive control). Panels (c) and (d) show the results of thermal lysis on paper at a lower temperature (60 °C) and reduced durations (5 min and 10 min). There are a few isolated colonies in these plates. These images were taken after incubation at 37 °C for 72 h. The large white spots at the centre of the plates are the paper discs.

### 2. Thermal stability of PicoGreen at 60 °C

In order to integrate fluorescence detection with the combined lysis and amplification step, we tested the thermal stability of PicoGreen at 60 °C under two different test conditions. In the first case, the dye was added to the sample prior to performing the reaction. In the second case, the dye was added to the paper substrate after the completion of the reaction. In both cases, the combined reaction was carried out for 60 min. Figure S2 shows that Picogreen in case 2 gives much higher fluorescence intensity compared to case 1, suggesting that the dye loses its efficacy when it is heated. Henceforth, we decided to add the dye to the amplification mixture after the reaction.

**Figure S2.**
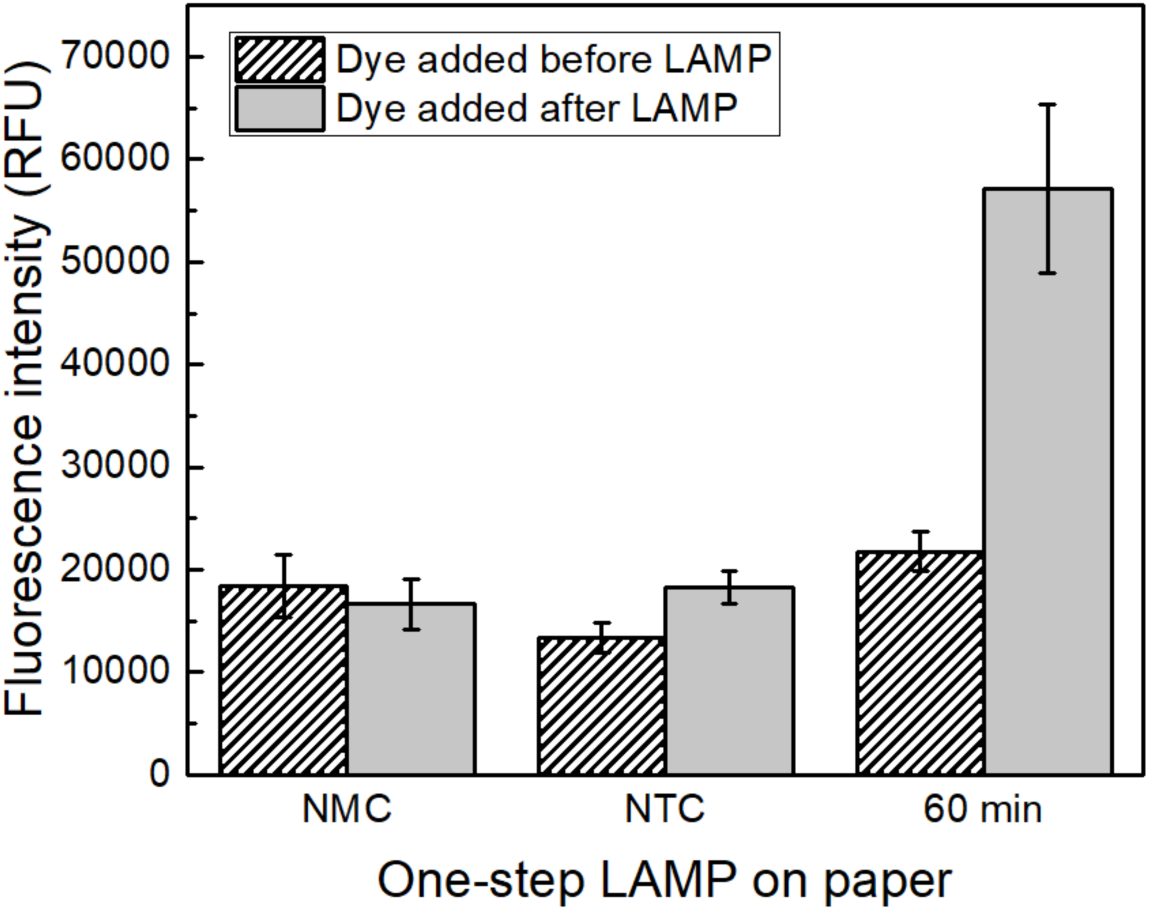
Thermal stability of the fluorescence marker PicoGreen. It was added to the reaction mixture before the reaction (case 1), as well as after the reaction (case 2). NMC indicates negative control without any mastermix in the reaction mixture. NTC indicates negative control without any bacterial cells in the reaction mixture. Lysis and LAMP were carried out at 60 °C for the recommended duration of 60 min. A large difference in the fluorescence intensities was seen between the two cases.

**Figure S3.**
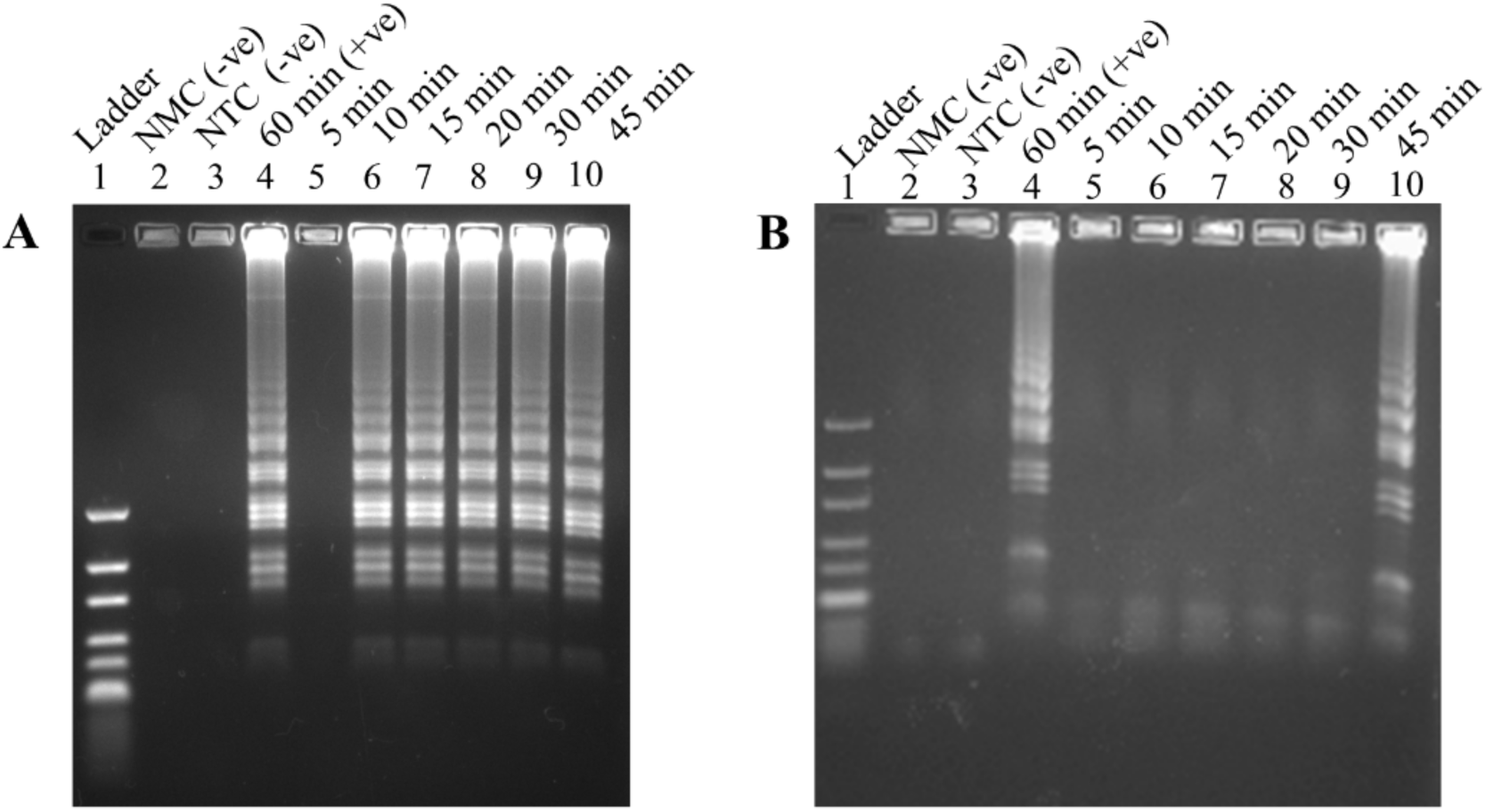
Estimation of the minimum reaction time for *E. coli* (panel A) and *M. smegmatis* (panel B) using gel electrophoresis. Lane 1: DNA ladder (10 bp to 300 bp). Lane 2: NMC indicates negative control without any mastermix in the reaction mixture. Lane 3: NTC indicates negative control without any bacterial cells in the reaction mixture. Lane 4: LAMP for the recommended duration of 60 min (positive control). Lanes 5-10: LAMP performed for 5 min, 10 min, 15 min, 20 min, 30 min and 45 min respectively. The minimum reaction times for *E. coli* and *M. smegmatis* are 10 min and 45 min respectively.

### 3. Estimation of the minimum amplification time using gel electrophoresis

We varied the total reaction times (lysis + amplification) from 45 min to 5 min for both *E. coli* MG1655 (10^7^ CFU/mL) and *M. smegmatis* mc^2^155 (10^8^ CFU/mL). The paper discs with the amplified DNA were loaded into agarose gels following the steps discussed in the Materials and Methods section. We found that gel electrophoresis could detect amplified DNA from *E. coli* after 10 min and from *M. smegmatis* after 45 min (figure S3).

### 4. Estimation of the limit of detection using gel electrophoresis

We tested the sensitivity of our protocol by taking several serial dilutions of *E. coli* and *M. smegmatis* cultures and performing the combined lysis and amplification assay. As shown in figure S4, using gel electrophoresis as the detection technique, our one-step amplification protocol can detect 1000 CFU/mL of *M. smegmatis* and 100 CFU/mL of *E. coli* (figure S4).

**Figure S4.**
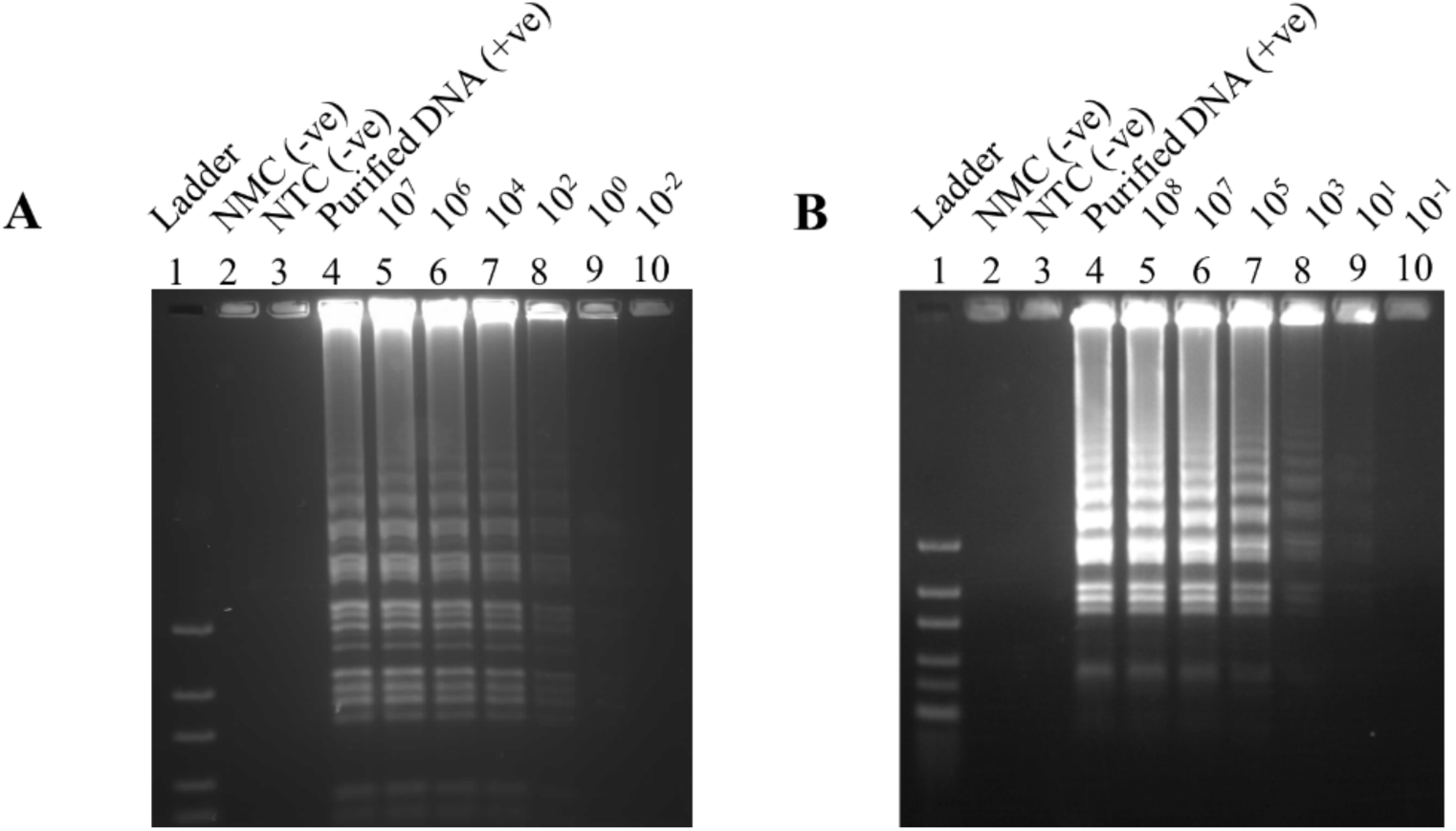
Detection sensitivity of our integrated assay with *E. coli* (panel A) and *M. smegmatis* (panel B) using gel electrophoresis. Lane 1: 10 bp to 300 bp DNA ladder. Lane 2: No master-mix control (NMC). Lane 3: No template control (NTC). Lane 4: LAMP of purified genomic DNA, used as a positive control. Lanes 5 – 10 in panel A: Reactions using different cell concentrations (10^7^ CFU/mL, 10^6^ CFU/mL, 10^4^ CFU/mL, 10^2^ CFU/mL, 10^0^ CFU/mL and 10^-2^ CFU/mL respectively). Lanes 5 – 10 in panel B: Reactions using different cell concentrations (10^8^ CFU/mL, 10^7^ CFU/mL, 10^5^ CFU/mL, 10^3^ CFU/mL, 10^1^ CFU/mL and 10^-1^ CFU/mL respectively). Gel electrophoresis can detect 1000 CFU/mL of *M smegmatis* and 100 CFU/mL of *E. coli*.

